# Higher rates of processed pseudogene acquisition in humans and three great apes revealed by long read assemblies

**DOI:** 10.1101/2020.06.07.139212

**Authors:** Xiaowen Feng, Heng Li

## Abstract

LINE-1 mediated retrotransposition of protein-coding mRNAs is an active process in modern humans for both germline and somatic genomes. Prior works that surveyed human data or human cohorts mostly relied on detecting discordant mappings of paired-end short reads, or assumed L1 hallmarks such as polyA tails and target site duplications. Moreover, there has been few genome-wide comparison between gene retrocopies in great apes and humans. In this study, we introduced a more sensitive and accurate approach to the discovery of processed pseudogene. Our method utilizes long read assemblies, and more importantly, is able to provide full retrocopy sequences as well as the neighboring sequences which are missed by short-read based methods reads. We provided an overview of novel gene retrocopies of 40 events (38 parent genes) in 20 human assemblies, a significantly higher discovery rate than previous reports (39 events of 36 parent genes out of 939 individuals). We also performed comprehensive analysis of lineage specific retrocopies in chimpanzee, gorilla and orangutan genomes.

## Introduction

Active human LINE-1s are responsible for various retrotransposons in the genome(Ostertag et al. 2003; Hancks and Kazazian 2012; Mandal et al. 2013), such as SVA, Alu and processed pseudogenes(Esnault et al. 2000), with a preference of L1 RNAs over non-L1 templates (Wei et al. 2001; Pavlicek 2002). While processed pseudogene formation usually leads to non-transcribed retrocopies because of the absence of promoter regions, in some cases, the retrocopies could encode proteins(Pink et al. 2011), regulate their parent genes(Cheetham et al. 2020) and ultimately possess significant functional implications such as carcinogenesis(Cooke et al. 2014; Poliseno et al. 2015). The mechanism behind parent gene preference hasn’t been fully revealed yet (Podlaha and Zhang 2009; Kazazian 2011; Richardson et al. 2015).

Processed pseudogene formation in the human genome has remained an active process both in germline and somatic, and non-reference events are described as gene retrocopy insertion polymorphisms (GRIPs). The term does not assume whether a given retrocopy is functional or not. We will also use ‘processed pseudogene’ and ‘gene retrocopy’ interchangeably without any implication about functionality. The total number of processed pseudogenes in the human genome has been estimated to range roughly from around 2000 to close to more than 10,000 and settled down on the higher numbers, depending on the criteria and discovery methods used (Zhang et al. 2003; Marques et al. 2005; Molineris et al. 2010; Frankish et al. 2019). Ewing et al. (Ewing et al. 2013) found 39 GRIPs representing 36 parent genes in 939 samples from 1000 Genome Project, and 26 GRIPs from 85 tumor-normal pairs from TCGA dataset, where the two sets overlapped for 17 GRIPs. Cooke et al. further examined 660 cancer samples, and found a total of 42 somatic events in 17 samples(Wei et al. 2001; Cooke et al. 2014).

Technologies such as Oxford Nanopore and Pacific Biosciences (PacBio) have enabled the sequencing inserts of tens of kilobases long, which could further be assembled into contigs of tens of megabases long. Given that 96% human transcripts annotated in Gencode are shorter than 10kb, the longest one being 109kb and the medium length being 2.9kb, we expect long read-based assemblies to reveal most processed pseudogenes. In this study, we introduced a novel processed pseudogene discovery approach which was more sensitive and accurate than SR-based methods, compared the findings with established results, and analyzed the L1 hallmarks as well as sequential landscapes around the retrocopies. Our results hinted that the GRIPs among the human population could be much more prevalent than previously suggested, lifting the rate from GRIPs of 39 events (36 parent genes) per 939 individuals to 40 events (38 parent genes) per 17 individuals. Moreover, we examined three great ape assemblies (chimpanzee, gorilla, orangutan) and provided an overview of their lineage specific events.

## Results

### Processed pseudogene polymorphism surveyed in 17 samples

We obtained 20 human assemblies (Seo et al. 2016; Vollger et al. 2019; Garg et al.). Each assembly approximately models a 3-gigabase human genome. We have more assemblies than samples because three samples (HG002, PGP1 and NA12878) have two assemblies per sample, representing the two phased haplotypes in a diploid human (see Methods). We aligned the contigs to GRCh38 and called structural variants (SVs) no shorter than 50bp. These SVs include both long insertions to and long deletions from GRCh38. On average, we called 11,101 long deletions and 13,969 long insertions from each assembly. We extracted the genomic sequences of protein coding genes (“gene reference”, including introns) using gene coordinates provided in Gencode and GRCh38, and splice-aligned the long SVs to the gene reference. We found an average of 7090 or 28% SVs in each assembly contained splicing signals. They were further screened for gene-like structures. We required that a retrocopy should contain at least 3 exons and few intron sequences. We then manually inspected the selected SVs to identify retrocopies (Figure 1B; Table S1; see Methods). We would from here refer to the retrocopies identified from long insertions/deletions as inserted/deleted retrocopies or such processed pseudogenes. Deleted retrocopies were the retrocopies that existed in the GRCh38 and not found in at least one of the compared assemblies, and inserted retrocopies were the ones found only in the assemblies but not the GRCh38. Human/ape lineage specific copies were also interpreted in this way (Figure 1A).

**Figure 1.**
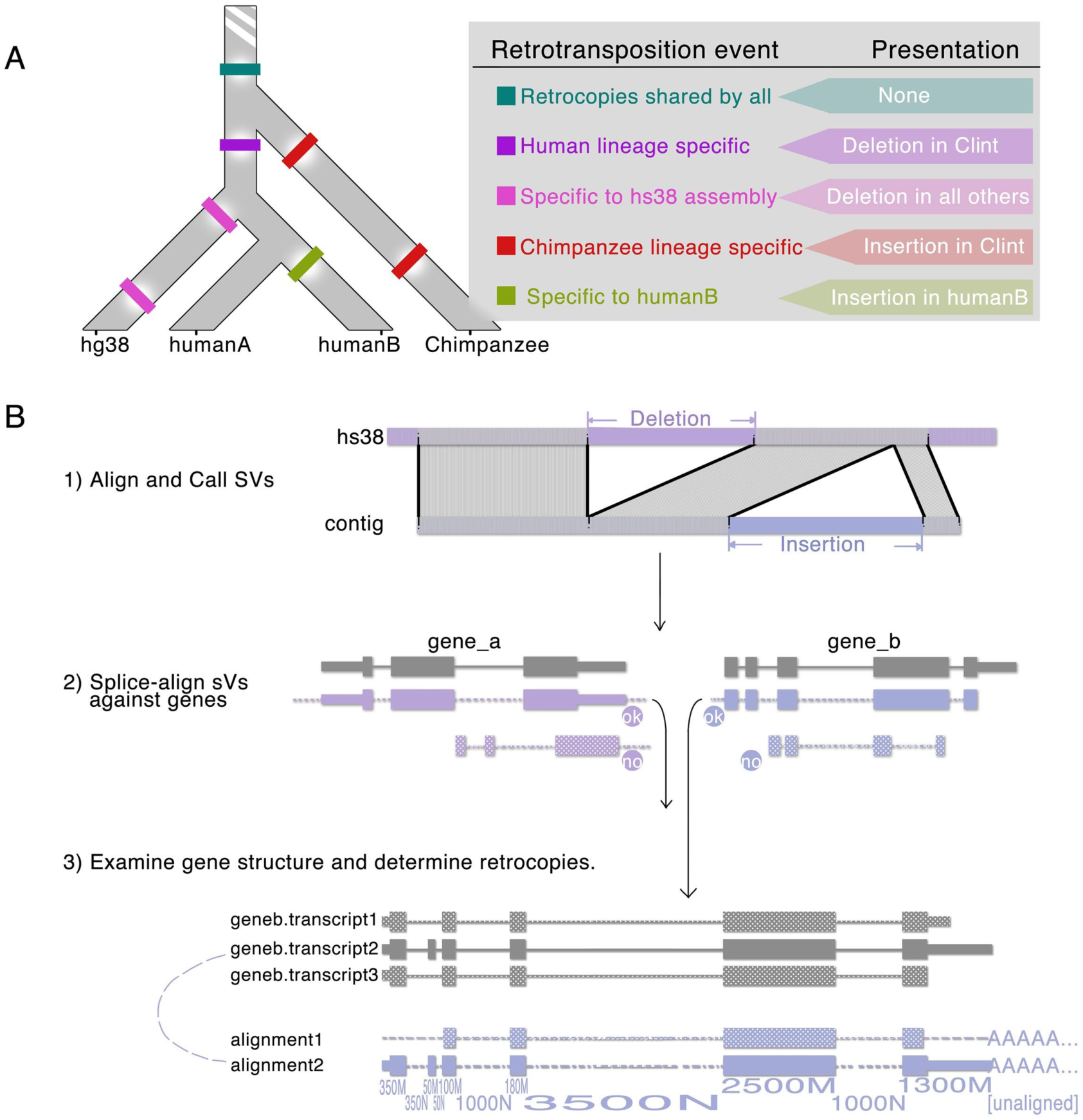
Overview of the gene retrocopy discovery approach. (A) Lineage-specific and non-reference retrotransposition events. For example, if an ancient event occurred before human-chimpanzee divergence, then theoretically its retrocopy wouldn’t be visible during structural variant calling since both GRCh38 and Clint would possess the copy at the same genomic location. On the other hand, if assemblyB recently gained a processed pseudogene, i.e. occurred after assemblyB’s ancestors diverged from GRCh38’s branch, then the retrocopy was expected to show up as a long inserted SV. (B) Diagram of detecting gene retrocopies from SVs.

We recognized a total of 176 inserted retrocopies derived and 148 deleted retrocopies derived from a total of 72 parent genes, where 38 inserted parent genes (AK2, AP3S1, C9orf85, CBX3, CIC, FAM91A1, GAPDH, GCSH, HNRNPC, IL6ST, KIAA2013, MOSMO, MTCH2, NANOGNB, NREP, NUDT4, PABPC1, PAIP1, PARP1, PDCL3, PPIA, PTGES3, RBMX, RPL10, RPL21, RPL22, RPL9, RPLP0, RPS26, RPS28, RPS3A, SKA3, SLC25A33, TDG, TERT, TYRO3, UPF3A, ZNF664) and 45 deleted parent genes (ABHD17A, AGGF1, ANKRD36C, ASNA1, ATXN1L, CIC, COX7B, DHFR, DYNLL1, EEF1A1, EIF4A1, FAM210B, FOXO1, GAPDH, GCSH, GNG10, HNRNPA1, ITGB1, KIAA2013, KRT18, MED15, RAMAC, RBM22, RHEB, RHOT1, RPL17, RPL21, RPL31, RPL36A, RPL41, RPL7, RPL9, RPS10, RPS15A, RPS16, RPS26, RPS28, RPS3A, RPSA, SLC25A33, SLC9A3, TOMM40, UPF3A, VOPP1, YBX3) were represented, respectively. Retrocopies of 11 parent genes (CIC, GAPDH, GCSH, KIAA2013, RPL21, RPL9, RPS26, RPS28, RPS3A, SLC25A33, UPF3A) were found in both deletions and insertions, implying that these genes were relatively more active in retrotranspositions (Figure 2; Table S5). Lengths of retrocopies varied from 157bp to 18328bp, with the mean value of 3068bp and the median of 1797bp. All deleted SVs except one have been unambiguously annotated as processed pseudogene for the GRCh38; the exception was region chr7:57410173-57418177, which was detected as deleted SV in PGP1 and HG03486. The current annotation of the region is “novel transcript” lncRNA AC237721.1.

**Figure 2.**
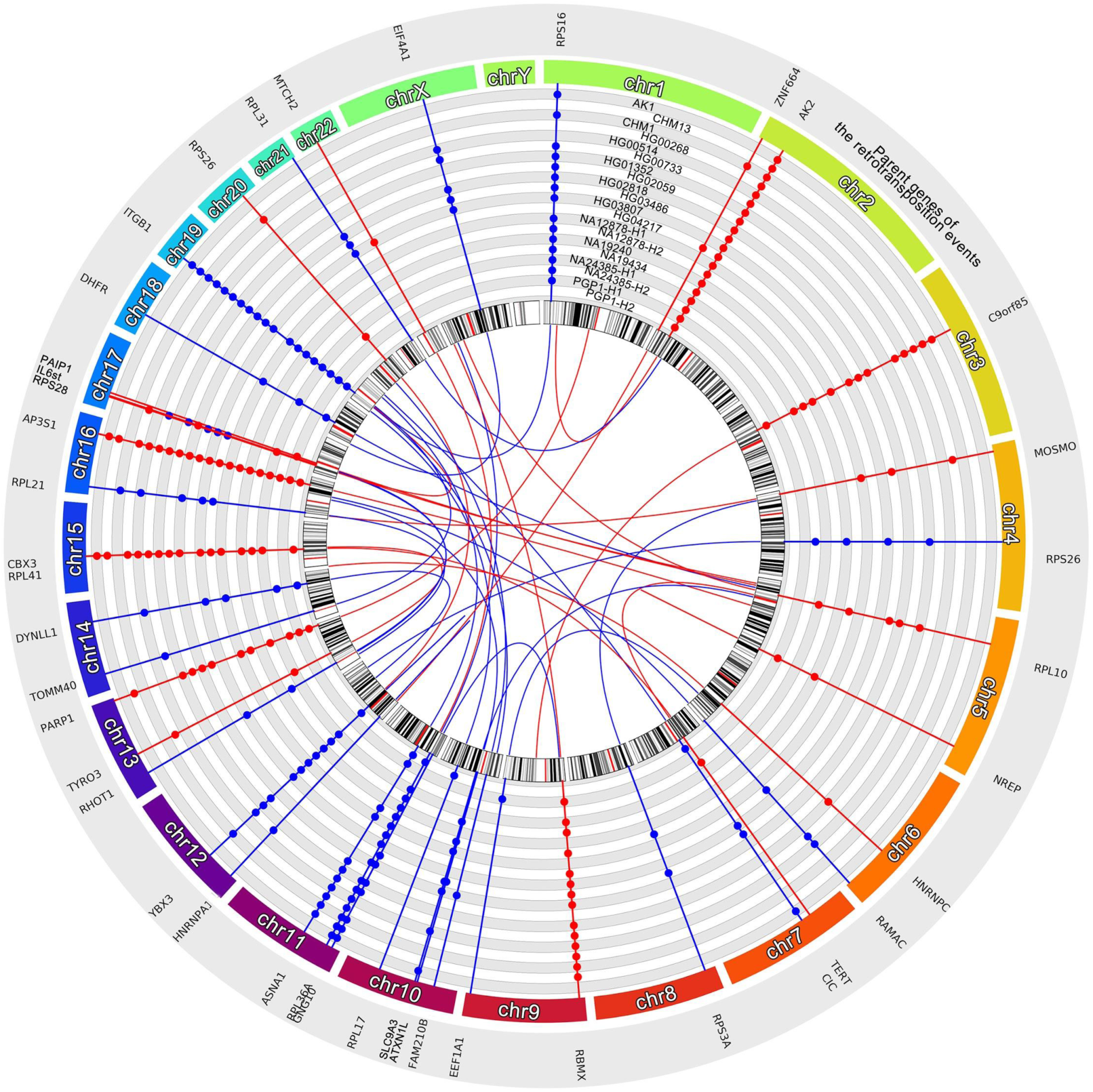
Chord diagram illustrating genomic locations of the gene retrocopies and the parent genes. The outer circles showed their presence in assemblies, where each gray or white lane represented one assembly. Blue lines and bars, deleted retrocopies. Red lines and bars, inserted retrocopies. Blue and red curves in the center illustrate the locations of parent genes (i.e. source of retrotransposition events) and the insertion point of the retrocopies.

With paired-end short reads, Ewing et al (2013) identified 6 novel retrocopies in HG00268 and 3 retrocopies in NA19434. We observed 6 of 9 events in our data. We are missing 3 previous findings probably because the two samples only have “collapsed” assemblies where one of the two parental alleles is randomly dropped. We were able to identify 8 more retrocopies in HG00268 and 7 more in NA19434. This demonstrates the enhanced power of long-read data.

To confirm the effect of collapsed assembly, we collected gene retrocopies from a collapsed assembly of NA12878 which was obtained independently from the two phased assemblies, and compared the findings. We expect the two phased assemblies to capture all retrocopies. We identified 14 heterozygous insertions/deletions from the two phased assemblies. However, only 5 of them are present in the collapsed NA12878 assembly. This suggests that using mostly collapsed assemblies, our approach underestimates the abundance of processed pseudogenes in these samples. Detecting pseudogenes from raw reads would address the issue. However, due to the technical difficulty in calling long SVs from reads, read-based discovery is challenging.

### Truncation and non-canonical alternative splicing were prevalent in human processed pseudogenes

PolyA tails were the most prominent L1 hallmarks that processed pseudogenes bear. We confirmed that 64.2% inserted retrocopies and 50.0% deleted retrocopies exhibited polyA tails (Table S2). Retrocopies without polyA tails had obvious 3’ truncation, either costing at least one exon(s), or a significant portion of the last exon (see below). The tails were relatively short compared to the polyA tail lengths in mature mRNAs which centered at 50bp(Chang et al. 2014), seldom exceeded 30bp and could be as short as 9bp. The polyA tails also harbored mutations or indels.

Since L1-mediated retrotransposition is target-primed reverse transcription (TPRT)(Cost et al. 2002), target site duplications (TSDs) were expected to be found at both ends of the retrocopies. We recognized a total of 61 unique TSD motifs in 59.6% inserted retrocopies and 71.6% deleted retrocopies (Table S2), with lengths ranging from 7bp to 40bp. In a few cases, TSDs also contained extra bases between the target site and the ends of retrocopies that measured as short as a single base and could go up to more than 20bp. While TSDs were expected to locate right next to the retrocopies, we still searched flankings up to 50bp for TSDs and manually inspected all sequences assisted by k-mer counting, and found no more convincing motifs. Compared to TSDs described for TYRO3, CBX3, GCSH, ZNF664, PPIA, RPL10 and RPS3A (Ewing et al. 2013), we confirmed identical or very similar TSDs. The mechanism or reason behind missing TSDs remained unclear, however.

Among both the inserted and deleted events, 32% lost the first or the first few exon(s) and 13% lost their last or last few exon(s). We also noticed that retrocopies of 6 genes (AGGF1, ASNA1, IL6ST, RPL10, SKA3, ZNF664) skipped one or more exons in the middle compared to the established transcript structures in Gencode. To verify whether the exon losses were a characteristic of these genes or were caused by non-canonical alternative splicings, we retrieved polished iso-seq data for validation from PacBio data release (https://www.pacb.com/blog/data-release-whole-human-transcriptome/) which provided whole genome full-length transcriptome for human brain, liver and heart. We also collected 8 iso-seq runs from SRP071928 (O’Grady et al. 2016), where the Burkitt lymphoma cell line Akata were profiled, as evidence supplementary to the PacBio data release. The iso-seq datasets were splice-aligned against the gene reference and ported to BED format for manual checks in IGV. Based on that, we determined that the inserted retrocopy of IL6ST might have lost exon 8∼13 from ENST00000502326 or ENST00000381298, since no such alternative splicing form was seen in iso-seq. Retrocopies of AGGF1 and ZNF664 lost the third exon and 5∼9 exons respectively. The fourth exon of ASNA1 lost a significant portion of sequence from the middle of it in the deleted retrocopy. Given that the neighbouring exons did not lose significant amounts of their content, we speculate that the missing exons were caused either by rare or tissue-specific alternative splicing, or by events during L1-mediated retrotranscription events instead of mutations afterwards.

Although 5’ truncation has been more frequently examined, sub-exon level 3’ truncation was also prevalent in retrocopies (Table S3), as already mentioned above about them and the polyA tail loss. Retrocopies of 38% parent genes experienced truncation at 3’ end that didn’t fully remove an exon. If only evaluating the mismatches and indels outside of the truncation or content loss events, 82% retrocopies had sequential divergences less than 10% compared to their parent genes, 67% had divergence scores less than 5%.

### The sequential landscape near processed pseudogene insertion point and the characteristics of parent genes

We found that the insertion sites of our identified events appeared to have no preference toward either genes or intergenic regions, and showed no preference toward coding strand or non-coding strand. Target sites of 26.4% inserted and deleted events located inside protein coding or lncRNA genes, similar to previously reported rate of 26.62% (Ewing et al. 2013). No host SVs were effectively inserted into protein coding exons. Although ribosomal genes were more active in the retrotransposition process(Zhang et al. 2006; Zerbino et al. 2018), a substantial portion of GRIPs or deleted retrocopies were contributed by lower-expressed genes and more inactive parent genes. We collected gene-level, tissue specific expression profiles from GTEx in transcripts per million (TPM) (Table S4). The max TPM values of 45% parent genes were less than 100, and only 33% parent genes had average TPM values higher than 100. Some genes, such as NREP, despite having high tissue specificity (neurons), exhibited retrocopies. Moreover, many GRIPs originated from parent genes that were rather inactive in ancestral events, i.e. the known retrocopies in GRCh38 that were also shared by other assemblies (Figure 3A). Since retrotransposition events in somatic cells would not be inheritable, polymorphism of processed pseudogenes is arguably accumulated events that happened to reproductive cells or during early stages of embryo development.

**Figure 3.**
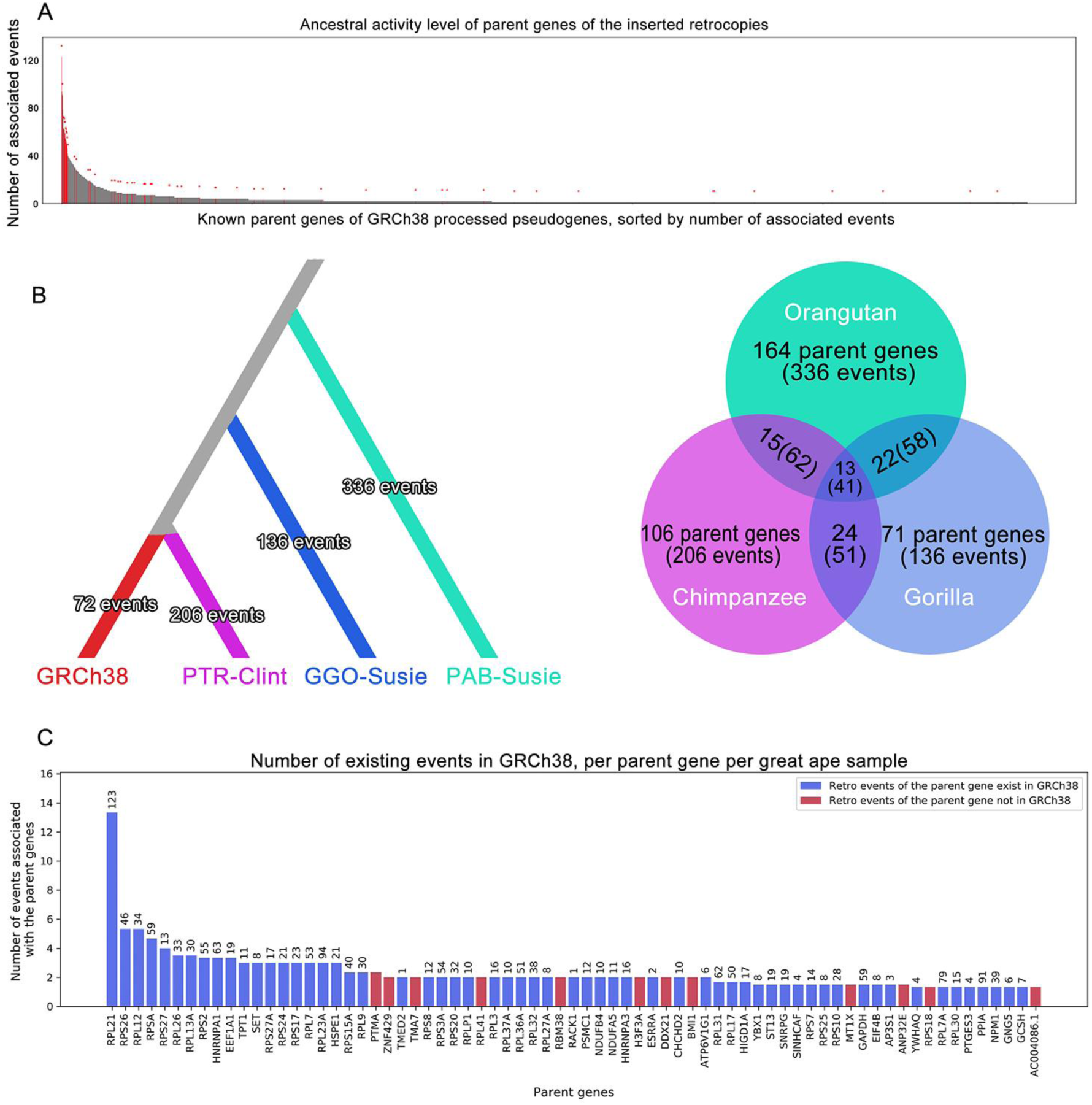
Parent gene active levels and lineage-specific events. (A) Recent GRIPs in the human population were not necessarily coming from the most active parent genes (e.g. ribosomal genes) in ancestral retrotransposition events. Left most bars illustrate parent genes with largest numbers of known retrocopies (Ensembl genome release 90, PseudoPipe). Red bars and arrows above them indicate GRIPs found in the 20 assemblies, gray bars represent other known parent genes and corresponding abundances of retrocopies. (B) Left: Processed pseudogene discovery in three great ape assemblies. Lineage-specific retrotransposition events of the human (represented by GRCh38), the chimpanzee (Clint), the gorilla (Susie) and the orangutan (Susie). Right: Number of retrotransposition events and corresponding parent genes unique to or shared by the great apes. (C) Genes active in retrotransposition in humans were not necessarily as-active in the apes. Heights of the bars show the counts of events in the apes; numbers above the bars indicate the counts of events in GRCh38.

### Lineage-specific processed pseudogenes in great apes revisited

We used three assemblies (chimpanzee, gorilla, and orangutan)(Gordon et al. 2016; Kronenberg et al. 2018) for processed pseudogene discovery in great apes (see Methods; Table S1). We recognized 127 deleted and 178 inserted retrocopies in the chimpanzee, 131 and 139 for the gorilla, 254 and 243 for the orangutan, as well as identifying the lineage-specific or shared events illustrated in Figure 3B and 3C (Table S7). Note that some parent genes shared by the apes might give rise to lineage-specific events. For example, non-reference retrocopies of PPIA were seen in the human samples and the three great apes, however all inserted to different spots (human at chr4, chimpanzee at chr6, gorilla at chr11 and orangutan at chr5). Retrotransposition activity level of gorilla, measured by lineage-specific Alus, has been reported to be lower than that of humans(McLain et al. 2013), possibly explaining the lower number of gene novel retrocopies seen in our gorilla sample; chimpanzee was reported to have significant higher speed of accumulating L1 insertions(Mathews et al. 2003), consistent with the observation here. Our results translate to roughly 46 gene retrotransposition events per million years in chimpanzees, 20/my for gorillas and 26/my for orangutans. Interestingly, the human had a rate of 16/my after human-chimpanzee divergence, based on our results, which is the lowest of the four.

The Chimpanzee Sequencing and Analysis Consortium reportedly found 246 lineage specific retrocopies corresponding to around 199 unique parent genes for the chimpanzee Clint (Chimpanzee Sequencing and Analysis Consortium 2005). We proceeded to compare the findings unique to chimpanzee lineage reported by the consortium and ours (Table S6; see Methods). 134 parent genes discovered were shared by both. 88 parent genes were found only by our method, being missed by the consortium because of either 1) the retrocopy presented in the assembly but failed to be recognized, 2) the retrocopy did not present in the assembly perhaps due to heterozygosity, or 3) the corresponding region not assembled. 65 parent genes were found only by the consortium. We failed to recognize these retrocopies due to either 1) the retrocopies were not lineage specific and not callable as SVs, 2) heterozygosity, or 3) the regions presented high divergence from the GRCh38, interrupting SV calling. Overall, we were able to reproduce most of the Chimpanzee Consortium’s results, and added more than 50% new parent gene discoveries.

We evaluated L1 hallmarks for the great apes using the same measurements we applied to human data, and yielded similar observations as in human retrocopies. 52% deleted SVs and 56% inserted SVs examined contained polyA tails up to 39bp and 54bp, respectively. 79% deleted retrocopies and 73% inserted retrocopies demonstrated clear TSDs. Truncations at the end of genes and losing part of or the entire exons in the middle were both prevalent, similar to the observations in human data. We define that losing more than 20% of an exon as truncation (if it’s the first or last exon) or content loss (if it’s not the first or last exon). Under this criteria, we found that 22% deleted retrocopies had 5’ truncation, 22% had 3’ truncation, and 12% experienced both; 32% inserted retrocopies had 5’ truncation, 28% had 3’ truncation, and 16% experienced both. Note that some cases described above might be non-canonical or lineage specific alternative splicing in the great apes.

NANOG is a DNA binding homeobox transcription factor that is involved in embryonic stem cell proliferation, renewal and pluripotency(O’Leary et al. 2016). Fairbanks et al. has reported that 9 out of the 10 processed pseudogenes of NANOG, except NANOGP8, were ancestral events and shared by humans and chimpanzees(Fairbanks and Maughan 2006). In addition to the NANOGP8 case, we found that human assemblies PGP1 (phased) and HG04217 (collapsed) gained novel retrocopies of NANOGNB at one same genomic location (chr1+:77406215-77406215) which was not found in ape samples, suggesting that the NANOG gene family is still relevant in L1-mediated retrotransposition events.

## Discussion

We described a long read assembly-based processed pseudogene discovery approach and showcased in 20 human and 3 great ape assemblies that the number of GRIPs in the population and retrocopies unique to the GRCh38 assembly could be much larger than previously reported, as well as expanding our knowledge of ape-specific processed pseudogenes. We provided a comprehensive overview of the insertion points, polyA tails, target site duplications and representation of parent genes’ exon structures by the retrocopy. This method also enabled us to examine the exact retrocopy sequences, which further suggested that cautious should be taken when interpreting sequence divergences between the retrocopies and parent genes among closely related species, like human and great apes. While collapsed assemblies could lose retrocopies that are discoverable by short read-based methods, we believe that as haplotype-aware assemblers quickly evolve, this concern would soon be resolved.

As a side note to Figure 2, the inserted retrocopies of AK2 (truncated, containing exon 4∼7 where exon4 lost its 5’ end), all originated from one same retrotransposition event, were presented in every examined assembly, which was not really explainable by incomplete lineage sorting. Through manual inspections in IGV, we speculated that GRCh38 might have misassembled around the insertion point (chr2:31823409), which is also the end point of the known processed pseudogene AK2P2 (truncated, containing exon 1∼4 of AK2), and caused AK2P2 to appear to be 3’ truncated as well as the illusion that other samples gained 5’ truncated AK2 retrocopies. The AK2P2 also had no polyA tail towards its end in GRCh38 (coordinates according to Gencode). Another countering theory would be, the GRCh38 contributor(s) originally had a full length AK2 retrocopy, but had lost the 3’ half due to other events before they were sequenced and correctly assembled to form the reference genome.

Interestingly, the insertion point (chr9:101358929) of PTGES3 retrocopies was too placed adjacent to a recognized processed pseudogene, AL359893.1. However, AL359893.1 showed no sequential similarity to PTGES3 and whose parent gene has been determined to be ZBTB, hinting that back-to-back processed pseudogene placement was not entirely impossible. We believe either way, the case of AK2 won’t change our main contribution and conclusions.

Our approach could also be utilized on long SV datasets without <u>long read-based assemblies,</u> although the yield would depend on SV calling quality and expect less discoveries than reported in this study. We tested on Hehir-Kwa et al.’s SV dataset which were obtained from 769 individuals of 250 Dutch families(Hehir-Kwa et al. 2016). A total of 20,494 long inserted SVs were selected and from which we identified 53 parent genes (<u>Table S8</u>). 16 of these 53 parents had neither novel inserted or deleted retrocopies observed in the humans and three great ape samples. As more long read datasets and de novo assemblies are becoming available in the near future, we believe that the divergence of processed pseudogenes could soon get better studied and cataloged.

## Materials and Methods

### Long read assembly-based processed pseudogene discovery

We obtained the following 21 human assemblies: AK1(GCA_001750385), CHM13(GCA_000983455), CHM1(GCA_001297185), HG00268(GCA_008065235), HG00514(GCA_002180035), HG00733(GCA_002208065), HG01352(GCA_002209525), HG02059(GCA_003070785), HG02818(GCA_003574075), HG03486(GCA_003086635), HG03807(GCA_003601015), HG04217(GCA_007821485), NA12878(GCA_002077035), NA19240(GCA_001524155), NA19434(GCA_002872155), haplotype-resolved assemblies of NA12878, NA24385 and PGP1 (ftp://ftp.dfci.harvard.edu/pub/hli/whdenovo/), and mainly used the following 3 assemblies of great apes: GCA_900006655.3 (Susie the gorilla), GCA_002880755.3 (Clint the chimpanzee) and GCA_002880775.3 (Susie the orangutan) represented three great apes. The assembly of kamilah the gorilla (GCA_008122165) was also processed and described in supplementary tables. We aligned the assemblies against hs38 with minimap2 (2.17-r974; -xasm5 -c --cs -z10000,200 for humans, -xasm20 -c --cs -z10000,200 for great apes), the results of which were then sorted (sort -k6,6 -k8,8n) and called for structural variants (SV) with minimap2’s paftools (k8 paftools.js call). We discarded SVs shorter than 50bp based on SV length distributions (empirical observations in this dataset) and exon length distribution of protein coding genes in the human genome(Gencode annotation). SV sequences were aligned against the genomic sequences of human protein coding genes (minimap2 - xsplice -c --cs -f10000 -N100 -p0.1; genomic sequences were defined by GRCh38 and Gencode V31), in search of properly spliced alignments which would be the implications of processed pseudogenes.

We defined the processed pseudogenes by the following criteria: 1) at least three exons of a multi-exon protein coding gene were partially or fully represented, 2) the best preserved exon should have at least 70% bases matched/mismatched, 3) no more than two introns were partially (>20%) or fully presented, and finally 4) manual inspections in IGV, since in corner cases, retained introns caused by non-canonical alternative splicing and non-retrocopy sequences aligned to introns could not be easily distinguished by only the above criteria. This was rare in both humans and the chimpanzee Clint, therefore the retrocopies found in great apes were not fully manually inspected due to the large amounts.

L1 hallmark PolyA tails were expected to be in the SVs, but not aligned to the gene reference. The immediate 50bp, or all contents left on the SV if less than 50bp, following the end of alignment blocks, were examined for polyA signals. The polyA tails were required to 1) appear close to the last alignment blocks, although allowing short insertions, and 2) contained at least 8bp of consecutive adenylate nucleotides (occasional mutations were allowed). Criteria 1 was enforced both by script and manual inspection. Similarly, target site duplications were searched at the flanking 50bp around both ends of the retrocopies, and their motifs were annotated by finding longest shared k-mers (k>=6). In cases where the polyA tail or TSDs were either too short or failed in manual inspection, they are labeled as “ambiguous”.

### Comparison with the previous chimpanzee retrocopy discovery

The Chimpanzee Sequencing and Analysis Consortium provided descriptions of the parent genes or their protein family instead of unique IDs or gene symbols, and a few descriptors differed only in one or two characters (e.g. a space or a dot). We manually curated the list, which yielded around 143 unique descriptors. Since these descriptors were still too broad for our comparison purposes (e.g. “Ras GTPase superfamily”), sequences of the 246 retrocopies were obtained and processed by our approach to be linked to specific parent genes, which linked them to 199 parent genes. Parent genes assigned to the retrocopies all matched their gene descriptors provided by the consortium. As we noticed that the consortium accepted retrocopies consisting of only 2 exons, we relaxed our discovery criteria to allow such candidates for both the consortium sequences and the chimpanzee sample accordingly, only in this part, to serve the comparison better. Lineage specific retrocopies reported in details still complied with the same criteria described above.

## Acknowledgement

This work was supported by the National Human Genome Research Institute (NHGRI) grant R01 HG010040, U01 HG010961 and U41 HG010972.

